# Systematic and quantitative analyses of pre-clinical *Mycobacterium avium* Lung Disease Tools for Drug Development and Transition from Animal Models to New Approach Methodologies

**DOI:** 10.1101/2025.07.10.664274

**Authors:** Moti Chapagain, Devyani Deshpande, Shashikant Srivastava, Tawanda Gumbo

## Abstract

Animal and new approach methodologies such as the hollow fiber system model [HFS] are used for *Mycobacterium avium-complex* [MAC] lung disease [LD] preclinical drug development. Our objective was to perform a systematic review to benchmark these pre-clinical tools. We performed a literature search to identify preclinical pharmacokinetics/pharmacodynamics [PK/PD] studies for MAC-LD. Preferred Reporting Items for Systematic Reviews and Meta-analyses was used for bias minimization. Twenty HFS-MAC and 3 mouse studies met PK/PD inclusion criteria. We created a novel quality score tool based on predictors of clinical response, design optimization, and information theory. The quality score was judged high in 10%, good in 50%, adequate in 30%, and poor in 10% of studies. Monte Carlo experiments for PK/PD target attainment were reported in 61% of studies. On repetitive sampling, the PK/PD target exposure estimate varied significantly between sampling days in 76% of studies. The solution was ordinary differential equations with parameter outputs such as γ [nonlinear kill-slope] and time-to-extinction applied to both HFS-MAC and patients’ sputa CFU/mL output. Next, we ranked the antibiotics by extent of microbial kill as fold-improvement over guideline-based therapy. The three top ranked drugs for microbial kill were omadacycline [69-fold], tedizolid [19-fold], and ceftriaxone [8-fold]. We recommend the HFS-MAC as tractable for exposure-effect and dose-fractionation studies, ranking antibiotics effect, and for translation to clinical doses. The analyses inform us of HFS-MAC recommendations for regulatory authorities and drug developers, including quality scores for optimal PK/PD design for target identification, resistance suppression, and choice of the best novel regimen.

---

*Mycobacterium avium complex* [MAC] is responsible for nearly 80% of all non-tuberculous mycobacteria [NTM] lung disease [LD] (1). Guideline-based therapy [GBT] includes a combination of macrolide [azithromycin or clarithromycin], ethambutol, and a rifamycin [rifampin or rifabutin] (2). Meta-analyses of clinical trials have demonstrated that GBT achieves 6-month sustained sputum culture conversion [SSCC] in only 43-53% of patients; the number needed to be treated is five to benefit one patient (3, 4). The predictors of patient clinical response to GBT are [1] pretreatment bacterial burden (*B_0_*), [2] drugs with high intracellular penetration such as macrolides and macrolide-based regimens, [3] presence of cavitary disease, and [4] cavities of >2cm diameter (2–5). Thus, therapy failure is due to pharmacokinetics [PK]-pharmacodynamics [PK] and the pathobiology of the disease. MAC-LD clinicopathological features include nodules and/or cavities, intracellular MAC in monocyte-lineage cells in granulomatous and necrotic lesions, with MAC *B_0_* median [range] of 5.17 [4.23-6.2] log_10_ CFU/mL in cavities and 3.0 [0.48-3.85] log_10_ CFU/mL in nodular/bronchiectatic lesions (6–9). The recent US Food and Drug Administration [FDA] and European Medicines Agencies [EMA] roadmaps state that new approach methodologies [NAMs] “offer the tools to assess safety, efficacy, and pharmacology of drugs and therapeutics *without* traditional animal models” and that such tools must be of “high relevance to human biology” (10, 11). Consistent with these recent regulatory agencies roadmaps, and MAC-LD clinicopathological factors associated with therapy failure, the hollow fiber system model [HFS] of MAC-LD [HFS-MAC] could be relevant to human biology as regards lung PKs, *B_0_*, intracellular nature of MAC, and drug diffusion properties (10–12). Here we performed a literature search for ALL preclinical PK/PD models, animal models and the HFS-MAC, to systematically characterize these models’ success and failures for benchmarking.

Antimicrobial PK/PD is a quantitative science of the relationship between drug PKs’ and antimicrobial effect such as bacterial [1] growth or [2] kill or (3) antimicrobial resistance [AMR] (13). PKs are captured by concentration-time profiles of drug and their shapes in lesions, as well as indices such as peak concentration [C_max_], 0-24h area under the concentration-time curve [AUC_0-24_], or % time concentration persists above MIC [%T_MIC_]. These are used to identify the PK/PD target exposure, which is the lowest exposure that kills the microbe and/or suppresses AMR, and the exposure that must be attained in the lung by a dose to cure MAC-LD. Ambrose *et al* have shown a strong correlation between the probability of PK/PD target attainment [PTA] of doses versus approval of new drug applications for antibiotic use in lung disease (14, 15). This PK/PD approach also improves the chances of success in treating patients in the clinic, while being cost and time effective, important for an orphan disease such as MAC-LD (15). Starting four decades ago PK/PD science was applied to Gram-negative bacilli and Gram-positive cocci, and then we applied to tuberculosis [TB] two decades ago, and more recently to MAC-LD and *Mycobacterium abscessus* LD (16–24). Here, we performed a systematic analysis for lessons learnt in MAC-LD preclinical PK/PD models.

## METHODS

### Rationale

Following work with the HFS model of TB [HFS-TB], which led to the qualification of the HFS-TB as a drug development tool by the EMA, we propose to achieve the same for preclinical models of MAC-LD (10, 11, 16–19, 25–27). Second, PK/PD work for MAC-LD has been ongoing for 15 years and has not been examined using meta-analytic approaches.

### Objectives and study question**s**

Objective 1: To develop a quality score tool to judge the quality of pre-clinical models for MAC-LD, for use by the academe, scientific journals, biopharma industry, and regulatory authorities.

Objective 2: To perform a systematic review of all pre-clinical models for MAC-LD that fulfil definitions of full PK/PD studies, for regulatory submission to the FDA and EMA.

Objective 3: To perform a quantitative analysis of all pre-clinical models for MAC-LD to rank new/repurposed drugs based on the extent of microbial kill, to design novel regimens.

### Definitions used as criteria for inclusion and exclusion of studies

The reason to perform antimicrobial PK/PD studies is to identify target exposures and dosing schedules. This is translated to optimal dosing regimens in the clinic, for maximal efficacy and minimal AMR. *In-silico dose* finding is most often implemented in Monte Carlo experiments, to account for PK variability in patients (28, 29). This approach integrates MIC distributions and generates PK/PD susceptibility breakpoints. These foundational concepts were used to define criteria for inclusion and exclusion of studies.

First, a PK/PD approach requires a PK system. This is because in patients both microbial kill and AMR are a consequence of fluctuating concentrations, the shape of the concentration-time curves, and periodicity [dosing schedule], not captured by static concentrations in time-kill curves (13, 16–19, 30, 31). In fact, the exact PK profile determines outcomes. As an example, in the HFS model of anthrax and in mice and macaques lung disease, the levofloxacin human half-life of 7.5h was compared to the mouse and rhesus macaque half-life of 2h at the same AUC (32, 33). The human PKs produced persistent reduction in bacterial burden while animal PKs led to failure and AMR (32, 33). Thus, static concentrations studies such as time-kill studies and stand-alone MIC results were defined as not fulfilling a PK/PD definition. Consequently, a preclinical model that did not report PKs from concentrations *measured* in the same experiment, did not fulfil the full PK/PD definition. An intended PK profile without measurements to back it up did not fulfil PK/PD criteria: “Trust but verify”.

Second, an exposure-response study must test several exposures to identify the PK/PD target exposure. So, what is the minimum number of exposures that define a full PK/PD study? The canonical model used for exposure-response analysis, the inhibitory sigmoid maximal effect [E_max_] or Hill equation, has the following formalism:

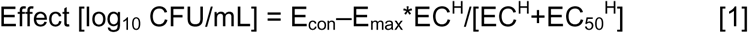

with the 4 parameters and [1] E_con_, [2] E_max_, (3) H, and (4) EC_50_ (24, 34, 35). EC_50_ or potency is the drug exposure mediating 50% of E_max_. If there are more model parameters [n] than data points [i.e., exposures or doses], the model will not have a unique solution and will be undetermined. Therefore, at a minimum n+1 or 5 exposure points are needed for optimal design of an exposure-response study. Moreover, based on both Shannon entropy and optimal Fisher information, exposures between those mediating 20% [EC_20_] and 80% [EC_80_] of E_max_, at least three exposures between of EC_20_ and EC_80_, are a minimum requirement for dose-fractionation studies (36). The definitions of at least five exposures for exposure-response study and three between EC_20_ and EC_80_ for dose-fractionation studies were used as inclusion criteria for PK/PD studies in the presented analysis.

Third, the same drug pair or triplets can have antagonism at some concentrations, additivity at others, and antagonism at yet other concentrations, at different sampling times (37–40). Therefore, for combination therapy, more than one dose needs to be examined in pre-clinical models. The PK/PD definitions of synergy or antagonism or additivity are most often based on two 2-parameter models and probability theory: Bliss independence and Loewe additivity. Bliss Independence has the following formalism:

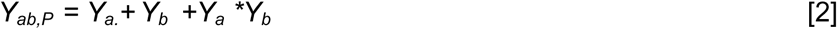

for interaction index, with *Y_a_* effect [i.e., inhibition or microbial kill] of drug “*a”* at specified dose, and *Y_b_*effect of drug “*b”* at specified dose. Therefore, there is need for at least three dosing regimens consisting of drug “*a”* alone, drug “*b”* alone, and “*a*b”* combined, plus non-treated controls [four regimens]. Exposures of each of the drugs to be examined should be at least three [EC_20_, EC_50_, and EC_80_] (41). Thus, for two drugs there is need of 16 combination pairs [including zero] at a minimum. Further, since interaction factor is defined based on the 95% confidence interval crossing zero, replicates are required. We defined combination therapy PK/PD study as being acceptable if there was minimum factorial design with at least the four regimens: *0, a, b, a*b*. These must have at least 2 replicates.

### Literature search strategy and study selection

We performed a literature search to identify PK/PD studies for MAC-LD, as defined by the criteria above. Stand-alone Monte Carlo experiments which used the PK/PD targets for dose selection were also included. The Medical Subject Heading [MeSH] terms and strategy used included either [1] “pharmacokinetics-pharmacodynamics” or “hollow fiber” OR “hollow fibre” or “mouse AND pharmacokinetics-pharmacodynamics” AND [2] either “*Mycobacterium avium”* or “non-tuberculous mycobacteria”. Bibliographies of original articles and guidelines were examined for additional relevant studies. Two authors [SS and TG] searched PubMed, Google Scholar, Inter-Science Conference on Antimicrobial Agents and Chemotherapy [ICAAC], ID week [the annual conference of the Infectious Diseases Society of America] and the yearly American Thoracic Society meeting, for studies published through March 31, 2025. When a conference abstract was found we also sought out the posters or oral presentations. If a conference presentation was published later in a scientific journal, the full publication was taken as the main reference. The two authors independently extracted the data into the prespecified table format. Consensus was reached for study inclusion after the discussion of each study. There was no exclusion of articles by language. Bias minimization was according to Preferred Reporting Items for Systematic Reviews and Meta-Analyses [PRISMA] (42).

### Data and outcomes collected

Data collected included number of units [animals, in vitro experiments units], replicates used, *B_0_* achieved in CFU/mL or CFU/lung, number of doses tested, type of study design [exposure-effect or dose fractionation or factorial design], number of bacterial isolates tested, and number of Monte Carlo experiments doses. The outcomes recorded were CFU/mL or CFU/lung on each sampling day, AMR population in CFU/mL, drug concentrations [mg/L or mg/g], PK/PD exposure values, and PK/PD parameters reported or linked to efficacy and AMR suppression. Calculated outcomes captured from the data or calculated by us were PK/PD target exposures, microbial kill below *B_0_,* EC_80_, and *in silico* dose-ranging to identify probability of target attainment [PTA] in published Monte Carlo experiments.

#### Quality criteria based on PK/PD science and predictors of clinical response to therapy

We proposed quality criteria based on experimental design optimization characteristics for the laboratory work, and on modeling the important predictors of patient response identified by clinicians, and “high relevance to human biology” (2–11). Since it is important to use multiple MAC isolates in order not to misestimate PK/PD targets, and since regulatory agencies guidelines state “that ∼4-5 organisms of the major target species or organism groups should be tested”, we also scored based on number of non-ATCC clinical isolates used (43). The proposed quality criteria were as detailed in **Supplementary Methods** and **Supplementary Table S1**. This initial tool was refined based on statistical analyses, to a final scoring tool.

### Data Synthesis and Analysis

The first level of analysis was qualitative. This was based on the main findings reported in each study. This approach is descriptive.

The second type of analysis was quantitative. For each study we captured the following data, and if data were not provided, we calculated the data:

1. Microbial kill below *B_0_* at E_max_ in CFU/mL and then ranked drugs by that metric. The microbial kill was weighted by the number of isolates used in the study, and fold difference with the three-drug GBT microbial kill among 5 isolates was calculated for normalization.
2. The target exposure, EC_80_.
3. Dose that achieves EC_80_ in lungs and the probability of toxicity at that dose.
4. Interaction index for combinations using factorial design.
5. The reliability of each study was qualified by the final quality score tool.

## RESULTS

### Selected PK/PD studies

We found 16 animal studies [37% of all studies identified], shown in the PRISMA flow diagram in **Figure 1** (44–59). Thirteen studies, shown in **Supplementary data Table S2** did not fulfill the PK/PD criteria for inclusion: most had less than 5 exposures tested, and all combination therapy studies excluded did not employ factorial design (44–55, 59). Three animal studies met the criteria, De *et al* exposure-effect and combination therapy [considered two studies], and Remal combination therapy study in BALB/c mice (56–58).

**Figure 1.**
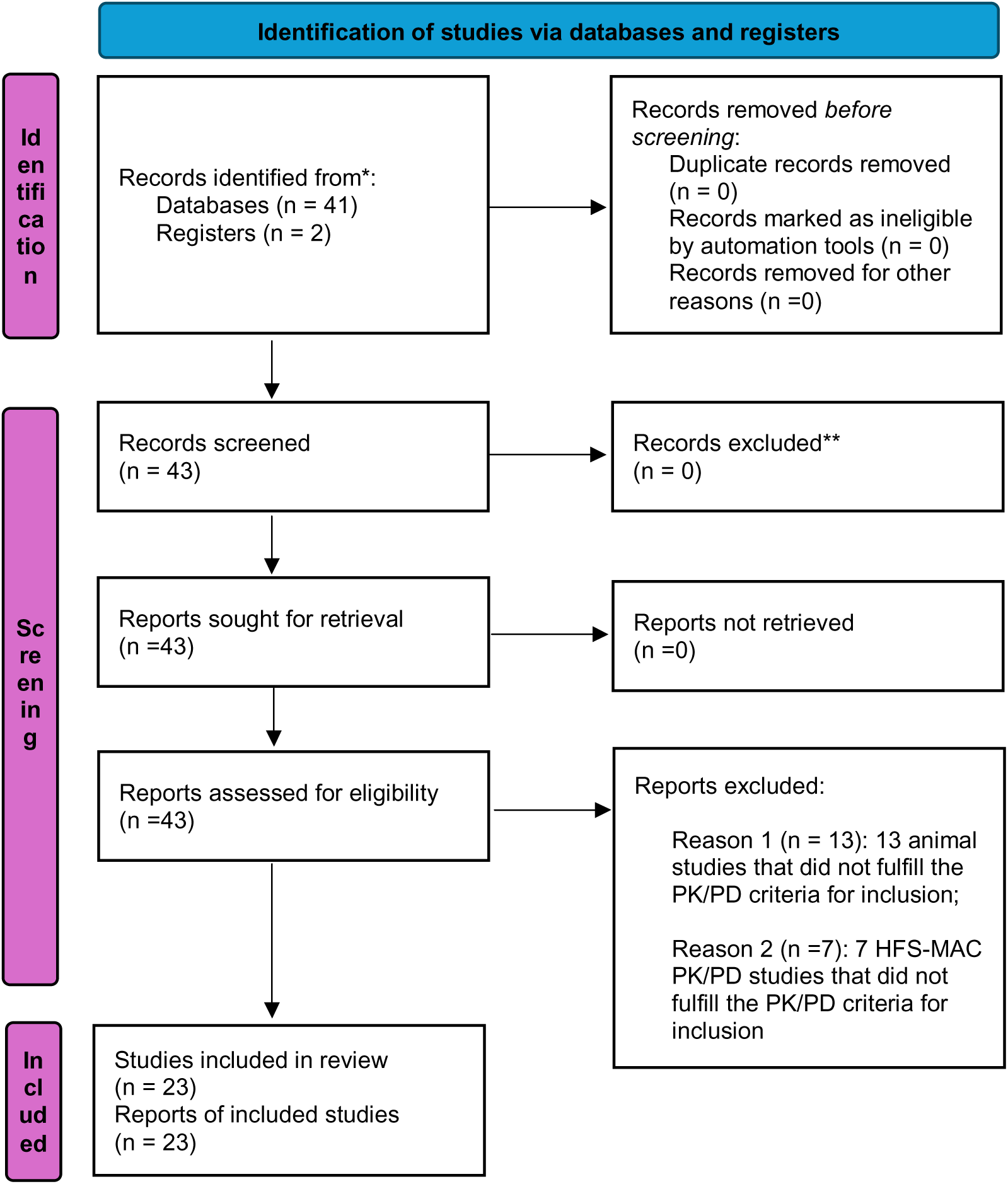
PRISMA Flow Diagram for Studies

We found 27 HFS-MAC PK/PD studies [63% of all studies identified], with first publication in 2010, shown in **Figure 1** (12, 60–85). Twenty HFS-MAC PK/PD studies and Monte Carlo experiments were met inclusion criteria (12, 60–78). Seven combination therapy studies were excluded because they did not meet the minimum definition of factorial design and are shown in **Supplementary Table S3** (79–85).

### Quality scores

The histogram of the quality scores using the initial tool is shown in **Figure 2A**. **Figure 2A** shows the distribution of scores including the combination therapy in HFS-MAC not otherwise discussed further because they did not meet PK/PD criteria and were included for tool development purposes only. The mean score for the excluded studies was 8.43 (95% Confidence interval (CI): 7.35-9.51) and would be categorized as poor quality or worse. **Figure 2B** shows the correlation coefficient between the different criteria was high [r>0.5 or R<-0.5] between “potency” [i.e., reports of EC_50_ and PK/PD target (potency)] versus doses [i.e., number of doses/exposures tested] criteria [Pearson r=0.91; p= 5.6 x10^-8^]. Thus, we removed the “potency” criteria from the tool. In addition, since all HFS-MAC studies basically scored the same for intracellular MAC criteria, and all mice had intracellular disease, this criterion was removed and enfolded into basic definition of an acceptable MAC-LD PK/PD study. The final criteria scoring system that was developed was as shown in **Table 1**. The total possible score was categorized as follows, [1] high score of >20, [2] good score of 15-20, [3] adequate score of 10-14, [4] poor quality score of 5-9, and [5] deficient and unreliable of less than 5.

**Figure 2.**
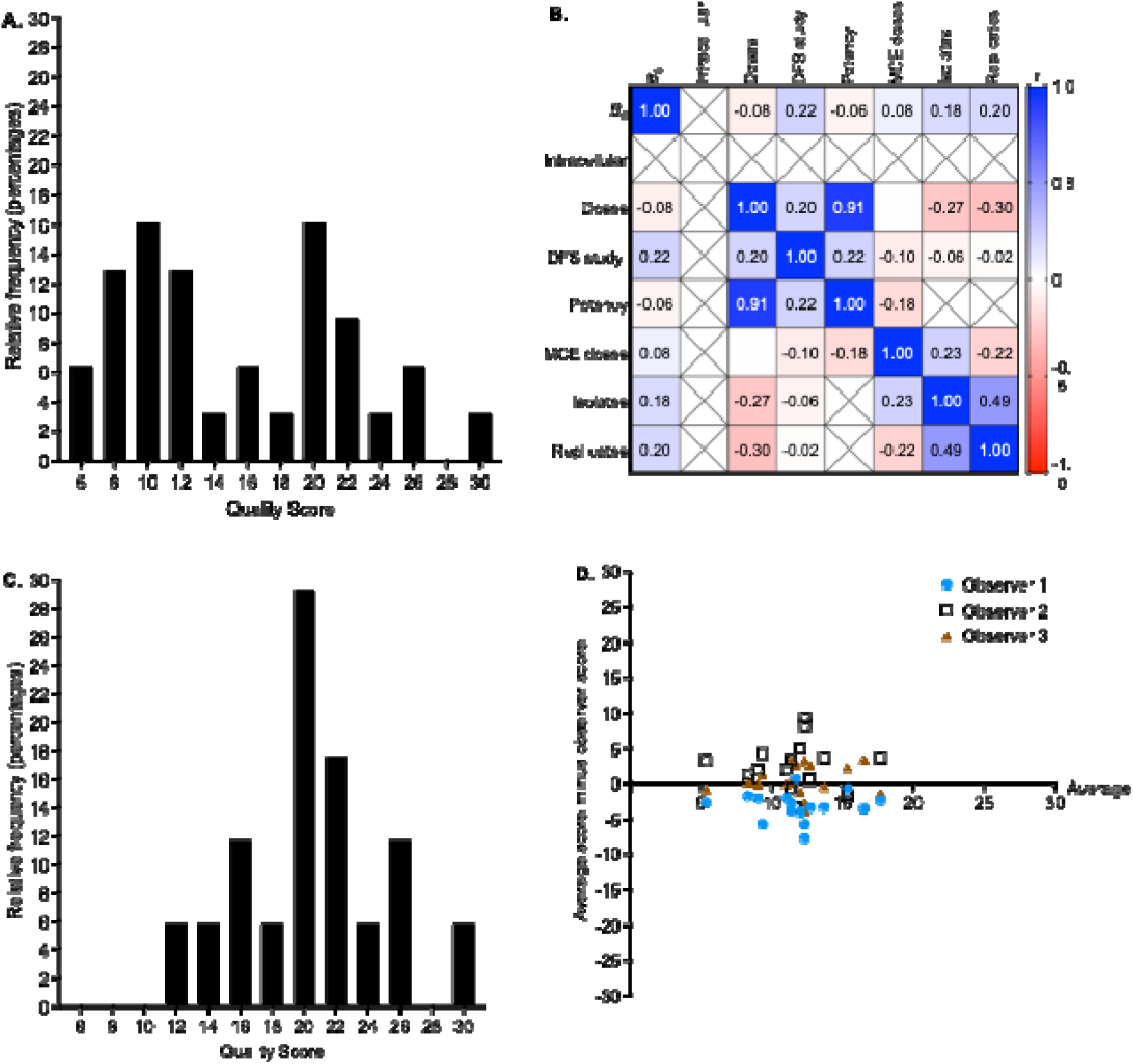
Hollow fiber system model of MAC Quality Scores **A.** Quality scores for all 43 studies. **B.** Correlation coefficients (Pearson) between different scoring criteria. **C.** Quality scoring with final tool, with only the studies that met inclusion criteria. **D**. Bland-Altman agreement analysis between the observers for the quality scores. The average score among all observers is shown on x-axis, while the difference between average and observer are on the Y-axis.

**Table 1.**
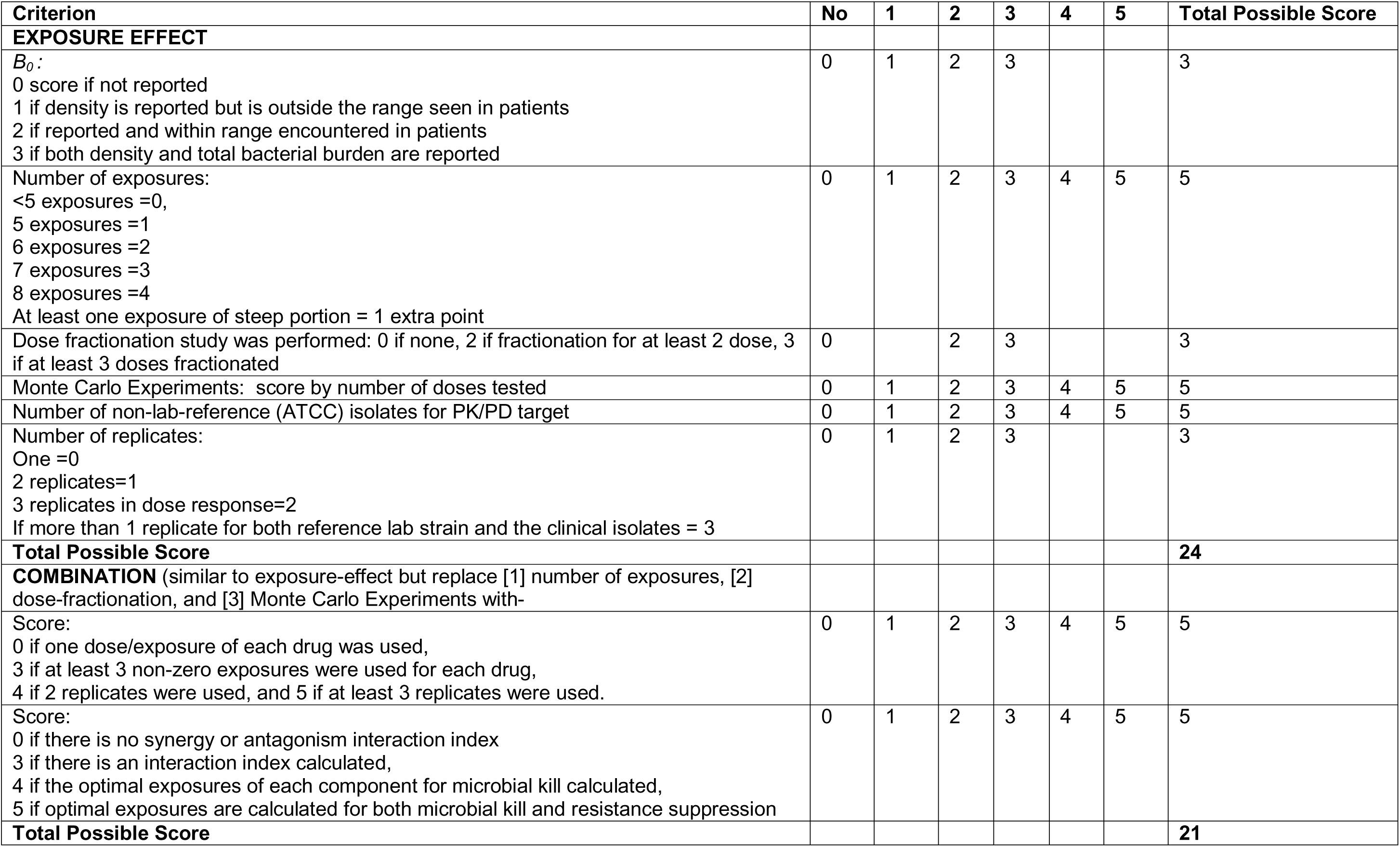
Final Proposed Quality Scores for preclinical pharmacokinetics/pharmacodynamics criteria

| Criterion | No | 1 | 2 | 3 | 4 | 5 | Total Possible Score |
| --- | --- | --- | --- | --- | --- | --- | --- |
| **EXPOSURE EFFECT** |  |  |  |  |  |  |  |
| *B<sub>0</sub>*:<br>0 score if not reported<br>1 if density is reported but is outside the range seen in patients<br>2 if reported and within range encountered in patients<br>3 if both density and total bacterial burden are reported | 0 | 1 | 2 | 3 |  |  | 3 |
| Number of exposures:<br><5 exposures =0,<br>5 exposures =1<br>6 exposures =2<br>7 exposures =3<br>8 exposures =4<br>At least one exposure of steep portion = 1 extra point | 0 | 1 | 2 | 3 | 4 | 5 | 5 |
| Dose fractionation study was performed: 0 if none, 2 if fractionation for at least 2 dose, 3 if at least 3 doses fractionated | 0 |  | 2 | 3 |  |  | 3 |
| Monte Carlo Experiments: score by number of doses tested | 0 | 1 | 2 | 3 | 4 | 5 | 5 |
| Number of non-lab-reference (ATCC) isolates for PK/PD target | 0 | 1 | 2 | 3 | 4 | 5 | 5 |
| Number of replicates:<br>One =0<br>2 replicates=1<br>3 replicates in dose response=2<br>If more than 1 replicate for both reference lab strain and the clinical isolates = 3 | 0 | 1 | 2 | 3 |  |  | 3 |
| **Total Possible Score** |  |  |  |  |  |  | **24** |
| **COMBINATION** (similar to exposure-effect but replace [1] number of exposures, [2] dose-fractionation, and [3] Monte Carlo Experiments with- |  |  |  |  |  |  |  |
| Score:<br>0 if one dose/exposure of each drug was used,<br>3 if at least 3 non-zero exposures were used for each drug,<br>4 if 2 replicates were used, and 5 if at least 3 replicates were used. | 0 | 1 | 2 | 3 | 4 | 5 | 5 |
| Score:<br>0 if there is no synergy or antagonism interaction index<br>3 if there is an interaction index calculated,<br>4 if the optimal exposures of each component for microbial kill calculated,<br>5 if optimal exposures are calculated for both microbial kill and resistance suppression | 0 | 1 | 2 | 3 | 4 | 5 | 5 |
| **Total Possible Score** |  |  |  |  |  |  | **21** |

The histogram of quality scores for this final tool is shown in **Figure 2C** for animal and HFS-MAC studies. The mean score for studies that met inclusion criteria was 15.35 (95%CI: 13.33-17.37). The quality score was judged high in 10%, good in 50%, adequate in 30%, and poor in 10% of studies. **Figure 2D** depicts a Bland-Altman agreement analysis between the observers for the quality scores. **Figure 2D** shows the average score minus observer’s score as an agreement metric; the overall average for the score was 11.28 [95% CI: 10.36-13.43] and the difference of scores between observers 0.03 [95% CI: -0.99 to 1.04]. The overall bias between all observers was 1.98 (95% confidence interval: 0.36-3.9), which is close to zero.

### Qualitative Systemic analysis findings

**Table 2** shows a qualitative analysis of the different studies. The characteristics of the drugs tested in these preclinical PK/PD models are shown in **Figure 3**. The most common pharmacophores studied were β-lactams [23.5%], followed by tetracyclines and oxazolidinones at 17.7% each, while macrolides were tested in only 11.8%. In the first three years, HFS-MAC studies lasted only seven days, but switched to 28 days starting 2012 leading to better mapping and PK/PD modeling of AMR sub-populations. All HFS-TB experiments underwent repetitive sampling and measured the drug concentrations versus time points that were achieved [which were all reported], for THP-1 cell count, as well as for both total bacterial burden and drug-resistant bacteria. No study reported the % coefficient of variance [%CV] between HFS-MAC replicates for drug concentrations or cell and CFU/mL counts. AMR universally arose to monotherapy and was concentration dependent.

**Table 2.** Qualitative Systematic Analyses Findings and Quality of Studies by Year of Publication.

| Ref. | Type of model | Year | Drug | Findings and conclusions | Quality Score | Study Quality |
| --- | --- | --- | --- | --- | --- | --- |
| (60) | HFS-MAC | 2010 | Ethambutol | 1. First HFS-MAC published, 7-day exposure-effect study<br>2. Studies of static concentrations of ethambutol with extracellular MAC over-estimate efficacy versus HFS-MAC<br>3. $C_{max}/MIC$ linked efficacy and resistance | 16 | Good |
| (61) | HFS-MAC | 2010 | Moxifloxacin | 1. 7-day exposure-effect study<br>2. Intracellular penetration overcomes protein binding<br>3. Proposed susceptibility breakpoint meant most clinical isolates resistant to moxifloxacin | 11 | Adequate |
| (12, 62) | HFS-MAC | 2012 | Azithromycin | 1. First 28-day exposure-effect study and 28 day dose fractionation study<br>2. The Antibiotic resistance arrow of time model proposed<br>3. Microbial kill and resistance linked to AUC/MIC, however on day 7 linked to % time above MIC ($\%T_{MIC}$) based on reading off the curves<br>4. Standard clinical doses sub-optimal | 16 | Good |
| | HFS-MAC | 2016 | Thioridazine | 1. Massive intracellular concentrations due to lysosomal trapping<br>2. $C_{max}/MIC$ linked efficacy and resistance based on day 7 results<br>3. Based on day 5 and day 14 results pattern was AUC/MIC driven microbial kill | 11 | Adequate |
|  | HFS-MAC | 2017 | Linezolid | 1. Linezolid bactericidal effect better than GBT components<br>2. In Monte Carlo experiments ~50% of clinical isolates resistant to linezolid | 17 | Good |
| (65) | HFS-MAC | 2017 | Tedizolid | Tedizolid killed more than $2.0 \log_{10} \text{cfu/mL}$ below $B_0$, highest seen yet | 10 | Adequate |
| (66) | HFS-MAC | 2017 | Ceftazidime/avibactam | 1. First β-lactam shown effective against MAC, at clinically achievable exposures<br>2. Ceftazidime was enhanced by avibactam<br>3. Efficacy higher than azithromycin and ethambutol. | 15 | Adequate |
|  | HFS-MAC | 2019 | Minocycline | 1. Minocycline is expected to be bactericidal at 400 mg/day<br>2. Use of Bayesian framework bacterial dynamics mathematical model based on competition model for drug-susceptible and resistant isolates | 13 | Adequate |
| (68) | HFS-MAC | 2020 | SPR720 | 1. PK/PD parameter linked to microbial kill was $AUC_{0-24}/MIC$<br>2. High intracellular penetration | 20 | High |
| (69, 70) | HFS-MAC | 2022 | Epetraborole (EBO) | 1. At the $EC_{80}$, EBO killed at least $2.0 \log_{10} \text{CFU/mL}$ below $B_0$<br>2. EBO + GBT demonstrated resistance suppression and faster microbial kill<br>3. Both microbial kill and AMR were $AUC_{0-24}/MIC$ linked | 24 | High |
| (56, 57) | C57BL/6 mice | 2022 | Epetraborole | 1. Epetraborole demonstrated potent in vivo efficacy against <u>5 MAC isolates</u> and significantly improved efficacy when combined with GBT<br>2. Epetraborole monotherapy better than GBT<br>3. No resistance emergence | 20 | Good |
| (56, 57) | C57BL/6 mice | 2022 | Epetraborole + GBT | More extensive microbial kill for epetraborole and GBT compared to either alone | 15 | Good |
| (71) | HFS-MAC | 2022 | Omadacycline | 1. Omadacycline concentration declines by 50% every 24h in MIC assay versus MAC doubling time of 16.3h, artefactually increasing MICs<br>2. γ-slope-based PD modeling more robust than day-to-day CFU/mL<br>3. Omadacycline more effective than GBT in same study | 14 | Adequate |
| (72) | HFS-MAC | 2024 | Sarecycline | 1. Due to high MICs and poor lung penetration $EC_{80}$ unachievable<br>2. "Paradoxical effect" observed with the $AUC_{0-24}/MIC$ of 285.39 exposure | 8 | Poor |
|  | HFS-MAC | 2024 | Ceftriaxone | 1. Exposure-response with reference MAC strain<br>2. GBT (azithromycin, ethambutol, rifabutin) killed 2 of 5 MAC clinical isolates, was partially successful in one, and failed in 2<br>3. Once-daily ceftriaxone monotherapy killed all for the 5 MAC below $B_0$<br>4. ODEs identified γ-slopes as PD parameter in 5 MAC strains<br>5. γ-slopes translated to SSCC in patients in 1,000 virtual patients, and ceftriaxone monotherapy predicted to shorten SSCC 2.35-fold | 20 | Good |
|  | HFS-MAC | 2024 | Ertapenem | 1. $E_{max}$ and $EC_{50}$ was better without avibactam than with avibactam<br>2. Ertapenem microbial kill below $B_0$ better than GBT drug monotherapies | 15 | Good |
|  | HFS-MAC | 2024 | Benzylpenicillin | 1. Better $EC_{50}$ without avibactam than with avibactam<br>2. Use of ODEs and γ-slopes that include model for resistance<br>3. Benzylpenicillin $\gamma=0.98$ (0.95–0.99) >azithromycin $\gamma=0.74$(0.65-0.86) and = azithromycin+ ethambutol $\gamma=0.94$ (0.82-0.99) | 16 | Good |
| (76) | HFS-MAC | 2025 | Tigecycline | 1. MICs of a drug not good indicators of potency or efficacy for MAC drug development because MAC is mostly intracellular<br>2. Static concentrations studies (e.g. time-kill) provide limited to no information as regards to PK/PD target exposures or efficacy<br>3. PK/PD target identified with ATCC reference strain was biased and imprecise<br>4. Inclusion of 5 clinical isolates gives a more robust target selection and estimate of $E_{max}$<br>5. Inhalation therapy was studied | 18 | Good |
|  | HFS-MAC | 2025 | Tedizolid | 1. Inclusion of 5 clinical isolates gives a more robust target selection that was 7 times higher than reference lab isolate (65)<br>2. 5 strains gave a more robust estimate of kill below $B_0$ that was very different from that derived with laboratory reference isolate (65) | 20 | Good |
|  | BALB/c mice | 2024 | Bedaquiline + clarithromycin | Bedaquiline plus clarithromycin combination therapy demonstrated enhanced efficacy compared to monotherapies | 8 | Poor |
| This study | HFS-MAC | 2025 | Clarithromycin | A total of 32 replicates, used to propose quality control criteria | NA | NA |

**Figure 3.**
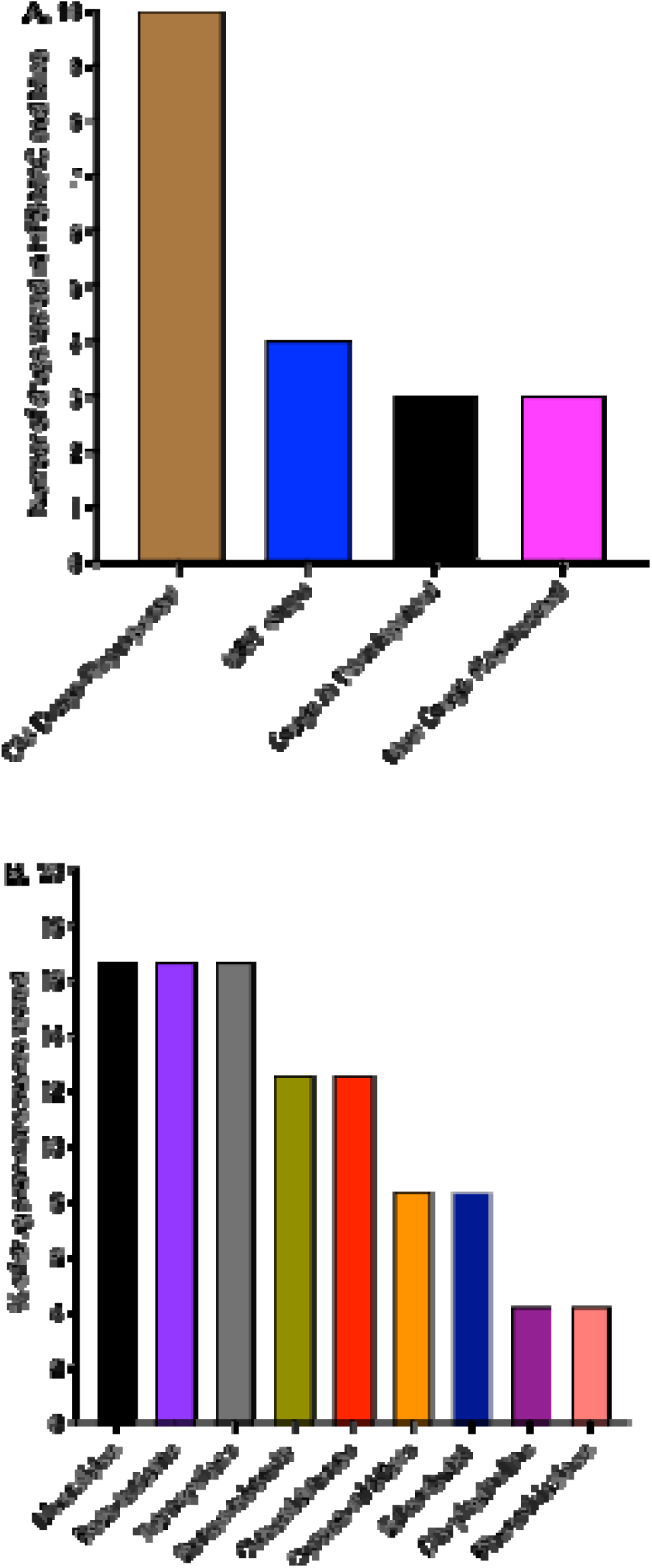
Drugs studied in HFS-MAC and murine MAC-LD studies **A.** Number of drugs in studies that fulfilled study acceptance criteria. **B.** Percentage of drugs studied, by pharmacophore.

In the first 10 years, CFU/mL output from HFS-MAC versus exposure on each sampling day was examined in the inhibitory sigmoid E_max_ and the antibiotic resistance arrow of time models, with parameter estimates used being those on the day with best Akaike information criteria (12, 60–66). However, for HFS-MAC studies that were longer than 7 days repetitive sampling revealed that in 13 out of 17 [76%] monotherapy studies, the EC_50_ and H varied between sampling days by a factor of 2 or more outside 95% confidence intervals, as could the PK/PD index [%T_MIC_ or AUC/MIC or C_max_/MIC] linked to effect (12, 63–67, 71–77). This presents problems as to which sampling day data to use to calculate the PK/PD target exposure.

Therefore, in 2019 sets of ordinary differential equations [ODEs] that gave parameter outputs that are vectors such as γ [non-linear kill slope), mutation rates [*m*], growth rates [*r*], and time to extinction [*τ*] were introduced for both HFS-MAC and patients sputa, with γ as new PD parameter (67, 71, 74, 75). Second, several studies demonstrated that the PK/PD target identification became more robust if up to five clinical MAC isolates were used in the PK/PD studies (65, 73, 76, 77). Commonly the reference ATCC MAC isolate mis-estimated PK/PD targets when compared to 5-isolate studies and to clinical studies (3, 4, 65, 73, 76, 77).

### Monte Carlo experiments

Fourteen of the 23 studies [61%] reported results of Monte Carlo experiments, for dose selection. These were reported mainly after monotherapy exposure-response studies.

### Quantitative analyses and ranking of drugs by extent of microbial kill below *B_0_*

Since the whole point of PK/PD studies is [1] identification of PK/PD targets in tandem with *in silico* based dose versus PTA in Monte Carlo experiments and [2] to find drugs with better efficacy [maximal microbial effect] than current GBT which performs poorly (3, 4), we performed a quantitative analysis that ranked drugs by how much better they compared to GBT [i.e., as fold of GBT]. **Table 3** ranks the microbial kill from the HFS-MAC studies and demonstrates that the top three ranked drugs, at doses that can be tolerated by people, were [1] 300 mg/day oral omadacycline or 35 mg/day inhaled tigecycline at 66-69-fold-GBT, [2] 200mg/day oral tedizolid at 19-fold-GBT, and 2g/day intravenous ceftriaxone at 8-fold-GBT. Based on Monte Carlo experiments to achieve PK/PD target exposure [PTA], the doses of each drug and the PK/PD susceptibility breakpoints shown in **Table 3**.

**Table 3.** Quantitative Systemic Analysis Findings and Ranking of Efficacy of Drugs for Monotherapy

| Drug [reference] | Weighted $E_{\max}$<br>($\log_{10}$ CFU/mL<br>kill below $B_0$) | Fold of 3-Drug GBT<br>Efficacy | PK/PD Target &<br>Dosing Schedule | Monte Carlo<br>Experiments-<br>generated Optimal<br>Dose (mg per day) | Route | PK/PD<br>breakpoint<br>(MIC; mg/L) | Predicted<br>Tolerability<br>of Dose |
| --- | --- | --- | --- | --- | --- | --- | --- |
| **MONOTHERAPY** |  |  |  |  |  |  |  |
| **HFS-MAC** |  |  |  |  |  |  |  |
| Thioridazine (63) | 4.68 | 227 | $C_{\max}/MIC=0.88$ | None | Oral | None | Poor |
| Omadacycline (71) | 4.16 | 68.63 | $AUC_{0-24}/MIC=9.9$ | 300 | Oral | 16 | Good |
| Tigecycline (76) | 4.14 | 65.576 | $AUC_{0-24}/MIC=33.65$ | 35 | Inhaled | | Good |
| Tedizolid (77) | 3.61 | 19.31 | $AUC_{0-24}/MIC=155.5$ | 200 | Oral | 8 | Good |
| Ceftriaxone (73) | 3.22 | 7.92 | $\%T_{MIC}=100\%$ | 2,000 | IV | | Good |
| Minocycline (67) | 2.9 | 3.77 | $AUC_{0-24}/MIC=63.1$ | 200 | Oral | 8 | Good |
| Moxifloxacin (61) | 2.30 | 0.95 | $AUC_{0-24}/MIC=163$ | 1,600 | Oral | 0.25 | Poor |
| Ceftazidime/avibactam (66) | 1.74 | 0.26 | $\%T_{MIC}=52\%$ | NA | NA | - | Good |
| SPR720 (68) | 1.54 | 0.17 | $AUC_{0-24}/MIC=4.09$ | 1,000 | Oral | 4 | Good |
| Ertapenem (74) | 1.47 | 0.14 | $\%T_{MIC}=100$ | 2,000 twice daily | IV | 4 | Good |
| Benzylpenicillin (75) | 1.4 | 0.12 | $\%T_{MIC}=84.6\%$ | 1,200 | IV | 64 | Good |
| Sarecycline (72) | 0.7 | 0.02 | $AUC_{0-24}/MIC=139.5$ | None | | None | Poor |
| Clarithromycin (this study) | 0.64 | 0.02 | - | - | Oral |  | Poor |
| Linezolid (64) | 0.38 | 0.01 | $AUC_{0-24}/MIC=43.7$ | 1,800 | Oral | 64 | Very poor |
| Azithromycin (12, 62) | 0 | <0.01 | $AUC_{0-24}/MIC=7.5$ | 8,000 | Oral | 16 | Poor |
| Ethambutol (60) | 0 | | $C_{\max}/MIC=1.23$ | 50 (mg/kg) | Oral | 2 | Poor |
| **MOUSE** |  |  |  |  |  |  |  |
| Epetraborole |  |  |  |  | Oral |  |  |
| **COMBINATION THERAPY** |  |  |  |  |  |  |  |
| 3-drug GBT in HFS-MAC<br>(73) | 211.0 | 1.0 |  |  | Oral |  | Poor |
| Epetraborole in mice |  |  |  |  | Oral |  |  |
| Bedaquiline + clarithromycin<br>in mice |  |  |  |  | Oral |  |  |

IV= intravenous

### Quality control HFS-MAC

For comparison and quality control purposes, we added a clarithromycin row in **Tables 2** and **3**, that is part of our unpublished internal controls for HFS-MAC over the last 15 years, given that clarithromycin is considered the primary drug driving outcome in MAC-LD. The data is from 32 HFS-MAC replicates [and 256 data points] of the American Type Culture Collection 700898 [ATCC] (12 HFS-MAC replicates) and five clinical isolates [20 HFS-MAC replicates]. The clarithromycin time-kill curves for the ATCC isolate versus the 5 clinical isolates that are part of our internal standards, are shown in **Figure 4A**, which demonstrates that the ATCC isolate gives an overoptimistic picture. **Figure 4B** shows the %CV between replicates on each sampling day; the higher %CV with ATCC is from work in earlier 10 years, while those in the recent 5 years had %CV below 20% between replicates on each sampling day. Overall, when all studies were considered, the %CV between replicates was 10.12% [95% CI: 7.01-13.24%]. Clarithromycin achieved 0.03-fold-GBT in ATCC isolate and 0.02-fold among 5 clinical isolates.

**Figure 4.**
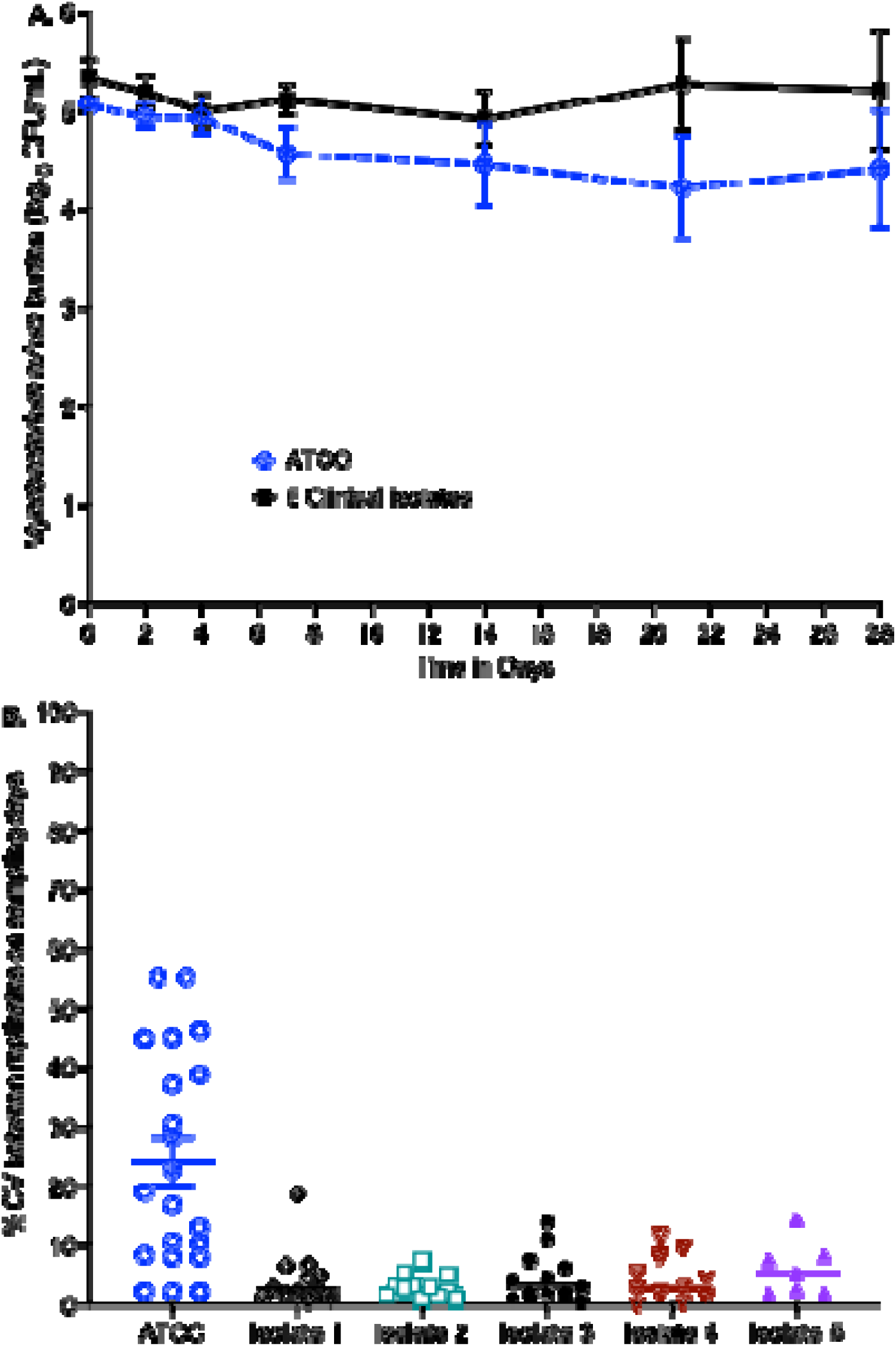
Clarithromycin effect in ATCC isolate versus a panel of 5 clinical isolates Symbols depict mean values. Error bars are standard deviation. **A.** Effect of clarithromycin on treatment with doses achieving EC_80_ in the lungs. **B.** Coefficient of variation of CFU/mL between replicates.

### DISCUSSION AND RECOMMENDATIONS

Sixty eight percent of MAC-LD PK/PD studies were performed in the HFS-MAC while 32% were in mice. Thus, the HFS-MAC has come a long way since the seven-day study with ethambutol 16 years ago (60). The HFS-MAC has become more sophisticated with repeated use, with longer duration of therapy in the system, which is now routinely over 28 days. The %CV between replicates has fallen from 24.24% in first 10 years to 4.53% recently. The HFS-MAC allows repetitive sampling and tracking of AMR evolution and microbial kill and their relationships to drug exposure. When the HFS-TB was qualified as a drug development tool, 22 HFS-TB studies had been published, a similar situation as with the HFS-MAC here (17–19). Our systematic analyses of both the HFS-MAC and murine models led to multiple recommendations.

### Quality scores [QS]

First, we developed a tool that scores pre-clinical laboratory models for quality and adequacy of the models, and for adequacy of PK/PD design. The final tool has 6 scoring criteria **[Table 1]**. We recommend this QS tool for HFS-MAC and mouse PK/PD work. The tool could also be used by academia for standardization and to assess rigor and reproducibility by the academe and granting institutes such as the National Institutes of Health. Indeed, the same concerns of rigor and reproducibility are often raised by editors and reviewers of scientific journals. For regulatory authorities [FDA or EMA], this tool could be used as minimal criteria for acceptance of PK/PD data biopharma for medical product development for MAC-LD. It could also be used to determine how much weight to give to a PK/PD result and dose choice for new drug applications. For biopharma companies, this could also be used as criteria to judge the quality of data from contract research organizations doing laboratory MAC-LD work.

### PK/PD design criteria for monotherapy and combination therapy for MAC-LD (5-5-5-25)

Based on pathological features that predict therapeutic outcomes, we recommend that the preclinical models of MAC-LD be intracellular and have the *B_0_* in the range observed in patient lesions (2, 6–9). Exposure-response PK/PD require at least <u>5</u> doses, and greater weight should be given to evidence from studies that have greater than 5 exposures. For combination therapy, since concentration-dependent synergy and antagonism are a fact of life, and since PK variability in patients will lead to situations in which each combination results *de facto* in a spectrum of exposures, at least three exposures should be tested in factorial design. To identify unbiased PK/PD targets, at least <u>5</u> MAC isolates should be examined in these studies (65, 73, 76, 77). Moreover, to account for biological variability, and to fulfil quality criteria scores, replicates should be used. We propose that a %CV between replicates on any sampling above <u>25</u>% be defined as a failure. We recommend documenting %CV between replicates for drug concentrations or cell and CFU/mL counts at each sampling time for all reports. We propose that the Monte Carlo experiments themselves examine at least <u>5</u> doses *in silico* for PTA to achieve target exposure and for probability of toxicity. Data generated from these more robust types of experimental design, and quality control, used to identify optimal exposures to be translated to human doses using Monte Carlo experiments, are crucial for better precision.

### The antibiotic resistance arrow of time

All PK/PD experiments should simultaneously identify and report the exposures associated with optimal kill and those associated with AMR suppression. In preclinical PK/PD studies, drug-resistant CFUs are captured by growing on agar supplemented with three times the MIC of a drug, in essence a three-fold increase in MIC, chosen to capture low-level resistance from efflux pump induction (12). Repetitive sampling in the HFS-MAC allows documentation of the evolution of efflux-pump induction and high-level chromosomal-mutation-related AMR versus drug exposure (12). On the other hand, the PK/PD susceptibility breakpoint identified in Monte Carlo experiment PTA defines resistance as the MIC above which patients fail therapy and is dependent on the dose used in the clinic and dosing route. Here in **Table 3**, we documented the PK/PD breakpoints for MAC-LD for drugs beyond clarithromycin. Clinical validation is still required. However, in the HFS-TB, such PK/PD breakpoints were identical to those picked by machine learning when clinical data was later examined (86–91).

### Recommendations of the best novel regimen for clinical trial testing

MAC-LD is an orphan disease. In addition, antibiotics have a negative net present value (92). This means that resources for developing new regimens are limited, as are the number of volunteers available for clinical trials. The most common approach has been to iteratively replace one of the drugs in GBT such as macrolides and rifamycins with repurposed ones (79–85). However, with this approach, it takes many years to develop a novel regimen and often can only examine single doses of each drug in combination (79–85). Moreover, this perpetuates the psychological “favorite child” based choice by academe: judging a drug’s utility based on being the discoverer or first adopter of the drug. We propose to select drugs for novel regimens-based ranking of how well they better than the combination GBT (macrolide plus ethambutol plus rifamycin). The top three ranked drugs in **Table 3** were omadacycline, tedizolid, and ceftriaxone: each surpassed GBT. We propose first performing HFS-MAC studies using factorial design for omadacycline, tedizolid, and ceftriaxone, with HFS-MAC output converted into ODEs, which can then be translated to SSCC in patients. A virtual 10,000 patient clinical trial can then be performed, followed by a clinical trial in patients that utilizes few patient volunteers.

### Limitations

First, there were few well designed mouse and combination therapy PK/PD studies. The large portion of studies excluded based on PK/PD quantitative criteria, which include three of our own, also scored poorly on our new quality score tool. There was no bias in our search, which means the poor scores are intrinsic to the studies. The most common reason what that the studies were not designed to identify target exposures in monotherapy or for combinations and could not inform on clinical doses or dosing schedules, which is the point of PK/PD studies. They missed the “5-5-5-25” criteria. Second, our novel quality score tool is yet to be tested by other investigators and thus remains a work in progress. Third, exposure-effect studies often did not include replicates. While technically the inhibitory sigmoid E_max_ is a regression that can handle lack of replicates, biological replicates are needed for better rigor and for quality control purposes. One of the recommendations resulting from current systemic analysis is to always have replicates. Fourth, use of multiple clinical isolates, at least 4, will be important for more robust estimation of target exposures. Both HFS-MAC and murine studies often had only the ATCC strains, with the limitation that results cannot be generalized. Fifth, while the role of biofilm in MAC lung is yet to be shown from human histology specimens, it cannot just be discounted. None of the models used explored that bacterial phenotype. Finally, there is always the inherent bias within ourselves given that we came up with the HFS-MAC model which is our own psychological “favorite child” over mice. That is the reason we introduced quantitative tools and criteria that are agnostic of models to address that bias. Our conclusions that the HFS-MAC model was more commonly used than mice in PK/PD studies, harmonizes with the FDA Roadmap statement: “*Over 90% of drugs that appear safe and effective in animals do not go on to receive FDA approval in humans predominantly due to safety and/or efficacy issues”* (10).

## REGISTRATION OF REVIEW

Not registered.

## ACKNOWLEDGEMENT.

None.

## DATA AVAILABILITY STATEMENT

The data for the results presented in the manuscript are publicly available.

## ETHICAL APPROVAL

Not applicable.

## AUTHOR CONTRIBUTIONS

Conceptualization and design, TG; Data analysis, MC, DD, TG and SS. TG wrote the first draft of the manuscript. All authors reviewed and approved the final version of the manuscript.

