## Supplemental Data for "Systematic and quantitative analyses of pre-clinical *Mycobacterium avium* Lung Disease Tools for Drug Development and Transition from Animal Models to New Approach Methodologies"

**SUPPLEMENTARY DATA**

**Supplementary Methods**

**Supplementary Tables**

### SUPPLEMENTARY METHODS

#### Quality criteria score development

First, we proposed the following quality criteria parameters:

1. Preclinical model must reflect the $B_0$ in patients in terms of density ($\log_{10}$ cfu/mL) and size of total bacterial burden: 0 score if not reported, 1 if density is reported but is outside the range seen in patients, 2 if reported and within range encountered in patients, 3 if both density and total bacterial burden are reported.
2. Intracellular infection model: 0 if its extracellular disease model, and 2 if intracellular
3. To construct the inhibitory sigmoid $E_{max}$ curve 5 exposures are needed at a minimum, and one must be on the steep portion of H. Frequently more are needed. A PK/PD study with less than 5 exposures scored a 0, that with 5 doses a 1, 6 doses a 2, 7 doses a 3, and 8 or more a 4.
4. A dose fractionation study was performed: 0 if none, 2 if done.
5. The potency ($EC_{50}$), and PK/PD target exposure ($EC_{80}$) and the confidence interval (or standard deviation) must be clearly stated as either AUC/MIC, $C_{max}$/MIC or $\%T_{MIC}$: if not score is 0, $EC_{50}$ alone 1, and $EC_{50}$ and $EC_{80}$ score of 2 – importance is dosing
6. MCE must be performed to identify best dose to achieve target exposure given PK variability, and an in-silico dose-response identified: score is 0 if no MCE, 1 if only one dose was examined, and 2 if at least 3 doses were examined.
7. Inclusion of non-reference lab isolates, 0 if none, and then 1 point each for the number of clinical isolates up to 5.
8. Number of replicates: score 0 if one, 1 if 2, 2 if 3 in dose response curves, and 5 if more than 1 replicate for both reference lab strain and the clinical isolates in #6.

The criteria are provided in tabulated form in **Supplementary Table S1**.

The total initial possible score was 30.

For combination therapy studies the scoring was as for exposure-effect studies, but we replaced criteria #3 to 6 as follows. A score is 0 if one dose/exposure of each drug was used, 3 if at least 3 non-zero doses were used for each drug, 4 if 2 replicates were used, and 5 if at least 3 replicates were used. Instead of $EC_{80}$ calculations, the score is 0 if there is no synergy or antagonism interaction index, 3 if there is an interaction index calculated, and 4 if the optimal exposures of each component for microbial kill are calculated, and 5 is optimal exposures are calculated for both microbial kill or slope and resistance suppression. The total score is 20 for factorial design. With the following categories: 16-20 is high quality, 10-15 is good, 7-10 is adequate, 5-7 is poor quality, and less than 5 is deficient and unreliable.

This initial approach was used and data gathered and scored. We then examined a correlation matrix to identify factors that had high r with other scoring criteria, as well as scores that were the same no matter which study was examined. Criteria with a Pearson $r>0.5$ correlation with other criteria were then removed, leaving one parameter. The final tool was then used to score the HFS-MAC and murine studies. The total possible score for exposure-effect PK/PD studies was divided into 5 categories in steps of 5 points: (1) high quality, (2) good, (3) adequate, (4) poor quality, and (5) deficient and unreliable.

#### SUPPLEMENTARY TABLES

**Supplementary Table S1. Initial Proposed Quality Scores for preclinical pharmacokinetics/pharmacodynamics criteria**

| Criterion | No | 1 | 2 | 3 | 4 | 5 | Total Possible Score |
|---|---|---|---|---|---|---|---|
| *$B_0$ :*<br>0 score if not reported<br>1 if density is reported but is outside the range seen in patients<br>2 if reported and within range encountered in patients<br>3 if both density and total bacterial burden are reported | 0 | 1 | 2 | 3 | | | 3 |
| Intracellular infection model:<br>0 if its extracellular disease model, and 2 if intracellular | 0 | | 2 | | | | 2 |
| Number of exposures:<br><5 exposures =0,<br>5 exposures =1<br>6 exposures =2<br>7 exposures =3<br>8 exposures =4<br>At least one exposure of steep portion = 1 extra point | 0 | 1 | 2 | 3 | 4 | 5 | 5 |
| Dose fractionation study was performed: 0 if non, 2 if done at least 1 dose, 3 if at least 3 doses fractionated | 0 | | 2 | 3 | | | 3 |
| Potency Reported: $EC_{50}$, $EC_{80}$ (or $EC_{90}$) exposure with CI or SE.<br>AUC/MIC, $C_{max}$/MIC or %$T_{MIC}$: if not score is 0, $EC_{50}$ alone 1, and $EC_{50}$ and $EC_{80}$ score of 2 | 0 | 1 | 2 | | | | 2 |
| MCE: score by number of doses tested | 0 | 1 | 2 | 3 | 4 | 5 | 5 |
| Number of non-lab-reference (ATCC) isolates for PK/PD target | 0 | 1 | 2 | 3 | 4 | 5 | 5 |
| Number of replicates:<br>One =0<br>2 replicates=1<br>3 replicates in dose response=2<br>If more than 1 replicate for both reference lab strain and the clinical isolates = 5 | 0 | 1 | 2 | 3 | 4 | 5 | 5 |
| **Total Possible Score** | | | | | | | **30** |

**Supplementary Table S2. Sixteen Animal Studies Versus PK/PD Inclusion Criteria**

| Author/reference | Year | Reason |
|---|---|---|
| Düzgüneş (1) | 1991 | 1. Single dose tested |
| Tomioka (2) | 1991 | 1. Less than 5 doses |
| Klemens (3) | 1991 | 1. Single doses tested<br>2. Combination had no factorial design |
| Tomioka (4) | | 1. Less than 5 doses |
| Klemens (5) | 1992 | 1. Less than 5 doses |
| Kailasam (6) | 1994 | 1. Less than 5 doses |
| Le Conte (7) | 1994 | 1. Less than 5 doses |
| Jagannath (8) | 1999 | 1. Less than 5 doses<br>2. No factorial design |
| Peters (9) | 2000 | 1. Less than 5 doses |
| Ellis(10) | 2014 | 1. Less than 5 doses |
| Grewal (11) | 2018 | 1. Less than 5 doses |
| Banaschewski (12) | 2019 | 1. Only 1 dose tested in PK/PD study |
| De (13, 14) | 2022 | **1. Included** |
| Rimal (15) | 2024 | **1. Included** |
| Cotroneo (16). | 2024 | 1. Only 3 doses examined<br>2. No drug concentration measurements<br>3. Combination not factorial design |

**Supplementary Table S3. Combination Therapy Studies That Failed the Minimum Definition of PK/PD Design**.

| Reference | Combinations | Score (total possible=18) |
|---|---|---|
| (17) | Azithromycin and ethambutol | 6 |
| (18) | Ceftazidime-avibactam, rifabutin, tedizolid, moxifloxacin | 9 |
| (19) | Thioridazine and moxifloxacin | 5 |
| (20) | Guideline-based therapy | 7 |
| (21) | Role of rifampin in Guideline-based therapy | 7 |
| (22) | Rifampin replaced by clofazimine | 9 |
| (23) | Minocycline replacing rifampin | 9 |
